# Top down information shapes lexical processing when listening to continuous speech

**DOI:** 10.1101/2022.05.31.494173

**Authors:** Laura Gwilliams, Alec Marantz, David Poeppel, Jean-Remi King

## Abstract

Speech is often structurally and semantically ambiguous. Here we study how the human brain uses sentence context to resolve lexical ambiguity. Twenty-one participants listened to spoken narratives while magneto-encephalography (MEG) was recorded. Stories were annotated for grammatical word class (noun, verb, adjective) under two hypothesised sources of information: ‘bottom-up’: the most common word class given the word’s phonology; ‘top-down’: the correct word class given the context. We trained a classifier on trials where the hypotheses matched (about 90%) and tested the classifier on trials where they mismatched. The classifier predicted top-down word class labels, and anti-correlated with bottom-up labels. Effects peaked ∼100ms after word onset over mid-frontal MEG sensors. Phonetic information was encoded in parallel, though peaking later (∼200ms). Our results support that during continuous speech processing, lexical representations are quickly built in a context-sensitive manner. We showcase multivariate analyses for teasing apart subtle representational distinctions from neural time series.

## 1 Introduction

Speech contains multiple sources of ambiguity. The same sounds, in the same order, can mean different things depending on context and expectations (Fodor et al., 1974; Simpson, 1984; Tabossi et al., 1987; Krovetz and Croft, 1992; Rodd, 2018). Despite the prevalence of ambiguity in speech (Rodd et al., 2002), listeners usually understand speech without difficulty and without error.

The saving grace of speech comprehension is context. Higher order structures of language, in the form of syntactic rules and overarching semantic topic, significantly constrain the space of plausible subsequent input (Miller and Isard, 1963; Spivey-Knowlton et al., 1993; Hagoort et al., 2004; Lee and Federmeier, 2009). Given the utterance ‘MJ looked at the stars using a…’, the upcoming word is more likely to be a noun, given the syntactic structure, and likely to be a word related to astronomy, given the semantic topic. This contextual guidance is particularly important in the case of lexical ambiguity, when a word (e.g., spell, watch, lean) has multiple meanings. Consider the difference between ‘Terri cast a spell’ and ‘Laura struggled to spell’, where the context favours one of the multiple possible interpretations of the ambiguous word.

The brain makes use of these constraints to guide interpretations of the input (Faust and Chiarello, 1998; Rodd et al., 2004; Gibson, 2006; Zekveld et al., 2006; Davis and Johnsrude, 2007; Cope et al., 2017). Violations of semantic or syntactic constraints are known to cause a reliable increase in brain responses, as measured with electro-encephalography (EEG) (Kutas and Hillyard, 1984; Friederici et al., 1993; Kaan et al., 2000; Hagoort et al., 2004; Gouvea et al., 2010), and sentences containing a lot of ambiguous words elicit stronger neural responses in the inferior temporal lobe and inferior frontal gyrus (Rodd et al., 2005), even when comprehension is equivalent to low ambiguity sentences.

How does the brain use contextual constraints to guide lexical access? Contextual influence has been most extensively studied in terms of how it disambiguates the semantics of a word. The exhaustive model posits that all possible meanings of a word are activated initially, weighted by their relative frequency of use (Duffy et al., 1988; Sereno et al., 2006; Gwilliams et al., 2017). Then, context serves to select the appropriate one (Simpson, 1984; Rodd et al., 2010; Simpson and Kang, 1994). Support for this model includes evidence from cross-modal priming studies, where a participant hears an ambiguous word such as ‘bug’, and after either a short or long delay is required to make a lexical decision on a written word that is related to one of the word’s meanings (e.g., ‘spy’ or ‘spider’). Even if the sentence context is heavily biased towards one meaning, participants’ short-delay lexical decisions are faster on words that shared either meaning with the auditory probe. In the long-delay condition, however, only the context-appropriate meaning is facilitated (Swinney et al., 1979; Onifer and Swinney, 1981). This supports the position that lexical processing initially activates all meanings of a word, which are then narrowed down to select just the context appropriate meaning.

While word meaning has been the primary focus of lexical ambiguity resolution, a word contains many properties, and semantics represents just a subset of them (Gwilliams, 2020). Words also contain grammatical features, such as word class (e.g., noun, verb, adjective) and the relevant properties of that grammatical class (e.g., number and gender, or tense and aspect). When ambiguity in word identity means uncertainty about word class (i.e., to pinV *versus* the pinN), the context also constrains the grammatical category of the word under scrutiny. It has been found that, contra to the semantic ambiguity literature, prior context serves to constrain the interpretation of speech input directly and immediately, such that only the appropriate word class is activated (Gaston and Marantz, 2018). This is more in line with context-dependent processing models, which posit that only the meaning consistent with the context is accessed, ignoring any other meanings that a word may assume in other contexts (Gennari et al., 2007; Mollo et al., 2018).

While humans constantly disambiguate speech using contextual information, when, where and how top-down grammatical disambiguation is implemented in the brain remain unknown. In the present study, we use lexical ambiguity to test two competing hypotheses. The first states that word class is initially generated bottom-up based on the phonological form of the utterance, and it is corrected later (if necessary) using top-down information. This is the order of computations that would be hypothesised based upon the semantic ambiguity resolution literature, in line with the ‘exhaustive model’. The second states that the top-down context guides word representations directly, without requiring an initial bottom-up parse. This is in line with the ‘context-dependent model’. For a visual schematic of the predicted results under each hypothesis, see Figure 1.

**Figure 1:**
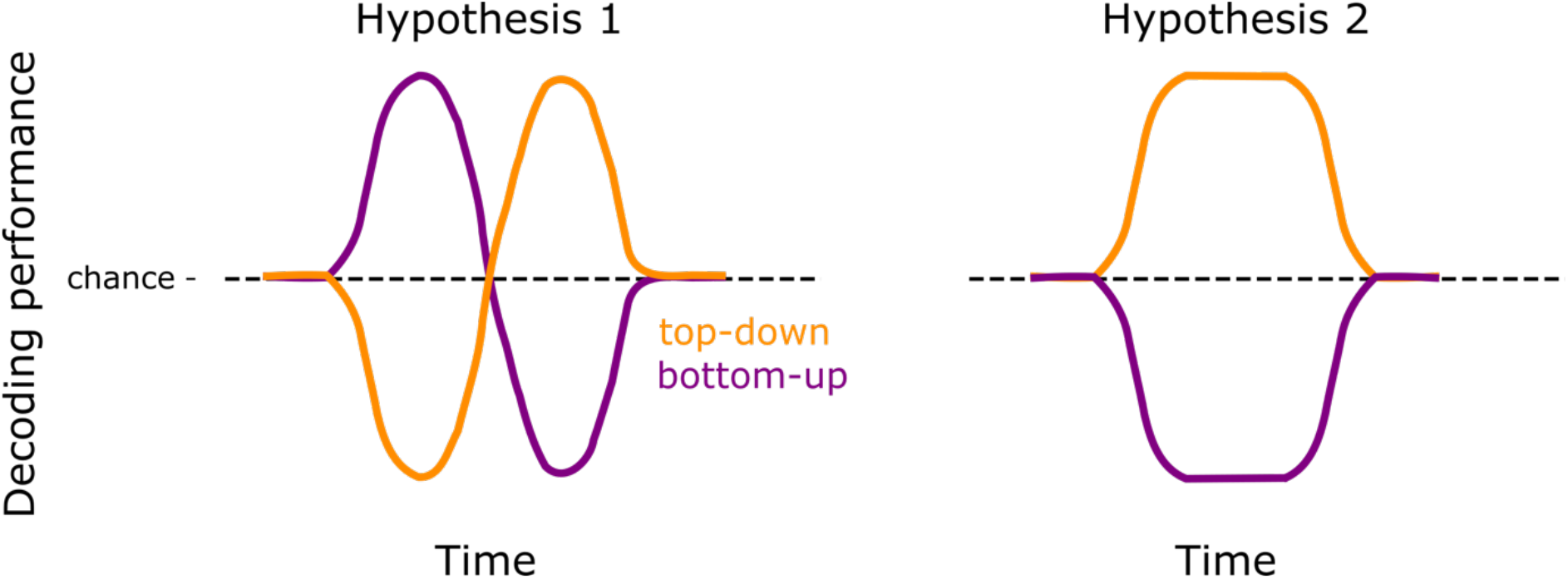
Analysis predictions. Schematic of expected results under our two hypotheses. The x-axis represents time relative to word onset. The y-axis represents decoding performance, where the dashed line is chance-level. The purple line represents the decoding of bottom-up labels of word class; the orange line represents decoding the top-down labels. The lines can go above and below chance performance: If the line goes below chance, it means that the classifier is systematically predicting a different class label. Hypothesis 1 predicts that evidence about word class from its phonological form is processed first (bottom-up information), followed by the higher-order sentence context (top-down information). Consequently, we would be able to first decode the bottom-up representation of word class, followed by the top-down representation. Hypothesis 2 predicts that top-down information exerts its influence on lexical representations directly. If this is true, we would expect to only be able to decode the top-down labels of word class from neural responses, with no trace of the bottom-up representation being encoded.

To adjudicate between these alternatives, we recorded magneto-encephalography (MEG) from 21 native English participants while they listened to four short stories. We modelled neural responses as a function of word class (e.g., noun, verb, adjective). Note that while we are not directly using a contextual model to predict one outcome or another, we reason that if the brain represents context-sensitive word class labels, this must have been derived using prior contextual information; and if the brain represents frequency-based labels, this must have been derived using learnt statistics over word forms. Within our ecological task of story listening, we found support for the second hypothesis: The pattern of activity that represents grammatical class in continuous speech directly encodes contextually sensitive grammatical class labels. The brain thus appears to make higher order interpretations accessible as early as possible to form a rapid and coherent understanding of speech in context.

## Methods

### 2.1 Participants

Twenty-one native English speakers were recruited for the study (13 female; age: M=24.8, SD=6.4). All were right-handed, with normal hearing and no history of neurological disorders. All provided their informed consent and were compensated for their time. The study was approved by the IRB committee at New York University Abu Dhabi, where the study was conducted.

### 2.2 Stimuli

Four fictional stories were selected from the Manually Annotated Sub-Corpus (MASC), which is a subset of the Open American National Corpus (Ide and Macleod, 2001) that has been annotated for its syntactic structure using Penn Treebank format.

We synthesised the stories using the Mac-OS text-to-speech application. Three synthetic voices were used (Ava, Samantha, Allison). Story duration ranged from 10-25 minutes. Participants answered a two-choice question via button press on the story content every ∼3 minutes. All participants performed this task at ceiling (98% correct).

### 2.3 Data acquisition

We used a 208-channel axial gradiometer MEG system (Kanazawa Institute of Technology, Kanazawa, Japan). Data were acquired at a sample rate of 1,000Hz, with online low-pass filter at 200Hz and a high-pass filter at 0.03 Hz.

Stimuli were presented to participants though plastic tube earphones placed in each ear (Aero Technologies), at a mean level of 70 dB SPL. Each recording session lasted about one hour. Each participant completed two recording sessions.

### 2.4 Pre-processing

We removed bad channels from the MEG data using an amplitude threshold cut-off of 3SD across all channels within a recording session, and linearly interpolated the bad channels using closest neighbours. We then applied a 1-50 Hz band-pass filter with firwin design (Gramfort et al., 2014) and downsampled the data to 250 Hz. The pre-processed continuous MEG data were epoched from -300 to 1,000 ms relative to word onset, and from -300 to 1,000 ms relative to word offset. No baseline correction was applied. All preprocessing was performed using the Python package mne, version 0.22.0.

### 2.5 Data annotation

We annotated the stories for the identity and timing of the 6,898 words they contained. We were primarily interested in two properties of these words. First is the word class predicted by the “bottom-up” hypothesis: i.e., the most frequent word class given its phonological form. We obtained these labels by querying the English Lexicon Project (Balota et al., 2007) for the word class of the words in our stories. The English Lexicon Project derives these labels from a collection of annotated spoken and written corpora. Second is the word class predicted by the “top-down” hypothesis, which refers to the word class that a word actually is assigned in the given sentence context. We obtained these labels from the manual Penn-Treebank syntactic annotation of our stories.

We focus on trials where the word class is an adjective, a noun or a verb in either the top-down or bottom-up definition of word class. This sub-selection yielded 3,941 epochs per subject per run, with 1698 unique words total, 1586 unique words where the two definitions matched and 155 unique words where the two definitions mismatched. The average word duration was 5.3 phonemes in the match condition and 5.6 phonemes in the mismatch condition.

Some of the words whose class differ across these two definitions were polysemous (had related meanings, and likely etymologically related) while others were homographs (had unrelated meanings, and likely not etymologically related). Unfortunately, we did not have enough trials to separate the analysis by this factor, but it would be an interesting avenue for future work to explore.

### 2.6 Analysis implementation

The majority of decoding analyses used the python package scikit-learn, version 0.24.1. This includes the functions LogisticRegressionCV, StandardScaler, and ShuffleSplit.

### Trial-type sets

We organised trials into two sets. “Match trials” refer to trials where the word class was identical across the bottom-up and top-down hypotheses. This encompassed 3,662 trials per subject, per run (93%). “Mismatch trials” refers to trials where the word class was different depending on how the word class was defined (279 trials, 7%). Our primary question was whether the bottom-up or top-down definition of word class best explains neural activity when the definitions conflict. For this, match trials were used to train the classifier, and mismatch trials were used to test the classifier.

In the mismatch trials, the word is used as its less frequent word class. For example, according to the Brown corpus, the word “watch” is used as a verb 86% of the time and as a noun 14% of the time. We consider “watch” a mismatch item when it is used as a noun, because its top-down usage (e.g., “the watch”) mismatches with its most common bottom-up label (verb).

Within the mismatch trials, the average token counts of the Brown corpus are 1102 for the top-down usage, and 1980 for the bottom-up usage. This means that the words are nearly twice as likely to be used as the bottom-up label than the top-down label.

### Optimisation

We used a logistic regression trained to perform a one-versus-all classification on the 3-class problem (noun, verb, adjective). The model was fit on each time sample independently, and no sliding window was used. We optimised the regularisation parameter at each time sample, selecting the best model fit on ten log-spaced alpha parameters from 1e-4 to 1e+4 (using LogisticRegressionCV).

We trained the classifer on the 3662 match trials and tested on the remaining 279 mismatch trials. We used the classifier to estimate the probabilistic class prediction for the held out test trials: i.e the soft-maxed distance of that trial from the hyper-plane distinguishing one word-class category from another.

### Evaluation

To evaluate decoding performance we compared the model’s probabilistic predictions (ypred) separately to the bottom-up labels (ybu) and to the top-down labels (ytd). We used the receiver operating characteristic (ROC) area under the curve (AUC) to summarise the likelihood that brain activity responded similarly to either hypothesis.

### Statistical assessment

To evaluate the reliability of decoding over time, we used a non-parametric temporal permutation cluster test across participants. First, we compute a t-value at each time-point by submitting the distribution of decoding accuracy across subjects to a one-sample t-test against chance performance. Second, we identified putative clusters by grouping consecutive t-values that exceeded a t > 1.96 (p < 0.05) threshold. Third, we compare the mean t-value within the cluster to a null distribution of t-values, which is formed by randomly flipping the sign of the distance from chance level, re-running the cluster forming step, and collecting the average t-value. This was performed 1,000 times. We consider clusters significant when their mean t-values exceeds 950 of the lowest values in the distribution (p < 0.05).

### Word-class decoding

To evaluate the base decoding of word class, we train the same decoder described above on just the match trials. The classifier was trained on 80% of trials and tested on the held-out 20% within a 5-fold cross validation loop. Performance was evaluated on the 5-fold average.

## 3 Results

Our goal was to understand the contribution of bottom-up and top-down computations during naturalistic listening, focusing on the lexical representation of word class (e.g., noun, verb, adjective). We defined word class in two different ways. First, a bottom-up definition: The most frequent word class ascribed to that phonological form (Figure 2A). For instance, the phoneme sequence ‘hammer’ is most often used as a noun, and so, this would be the bottom-up definition regardless of the context it was being used in. Second, a top-down definition: The true word class assigned given the sentence it is occurring within (Figure 2B). The top-down label of word class corresponds to when the word is being used as its less frequent word class (e.g., ‘hammer’ as a verb).

**Figure 2:**
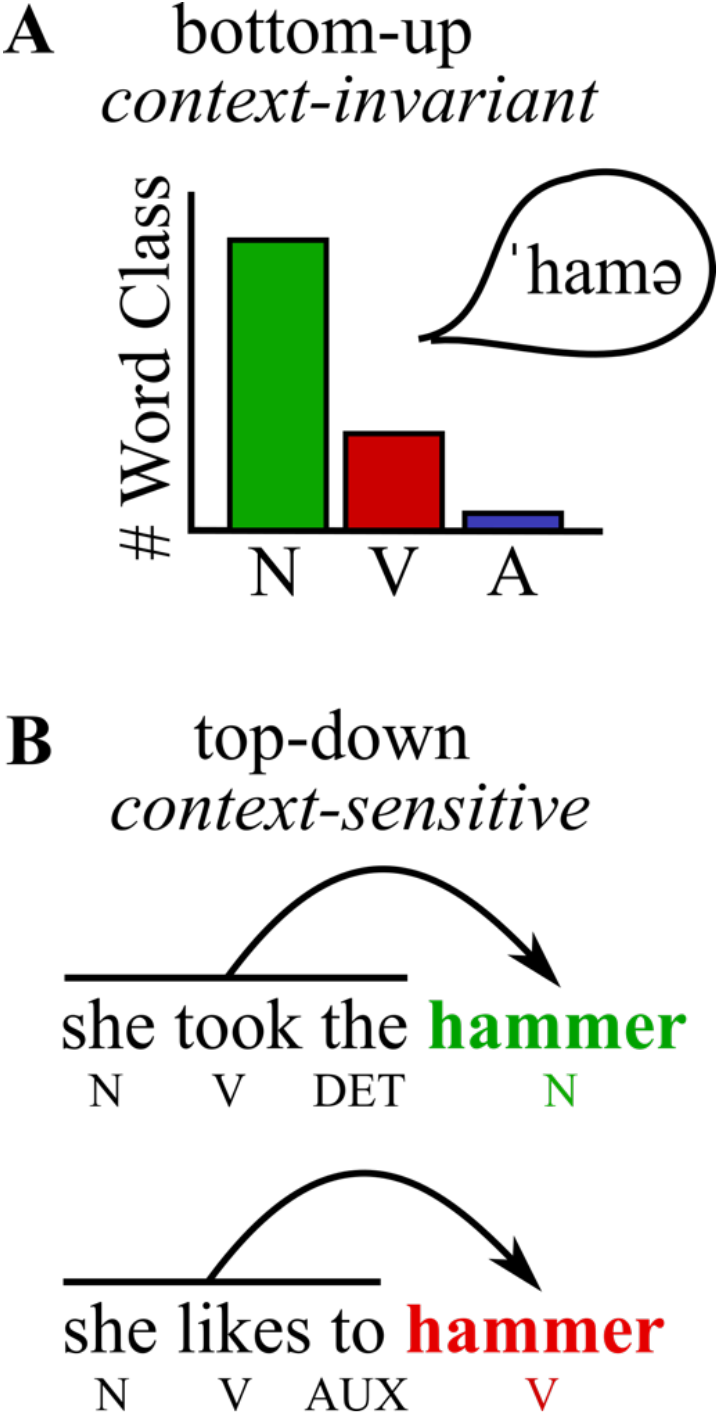
Definitions of word class. A: Bottom-up word class corresponds to the most frequent word class category assigned to a particular phonological form. In this case, the word ‘hammer’ is more often used as a noun than a verb; therefore, the bottom-up definition of word class for this word is noun. B: Top-down word class corresponds to the grammatical word class the item is being used in within the sentence. The two sentences provide examples where the same word, ‘hammer’ is being used as a noun (the sentence above) and as a verb (the sentence below).

First, we tested a hypothesis-free encoding of word class, on trials where top-down and bottom-up labels matched. We observed that word class was decodable from the neural responses to the spoken narratives. To test this, we subset all words that were either a noun, verb, or adjective, and whose bottom-up and top-down definitions gave the same word class label (3,662 trials). We then used logistic regression to distinguish the three classes, and the Area Under the Curve (AUC) to summarise decoding performance. Using a temporal permutation cluster test, we found that nouns were decodable during the entire epoch, timelocked both to word onset (average t = 8.1, p < .001) and word offset (average t = 9.2, p < .001). Verbs were decodable from -40 to 1050 ms from word onset (average t = 7.3, p < .001) and from -400 to 1080 ms relative to word offset (average t= 8.2, p < .001). Although adjectives were not significantly decodable relative to word onset after correction for multiple comparisons, they were decodable relative to word offset from 210-320 ms (average t = 3.6, p = .004) and from 370-810 ms (average t = 4.4, p < .001).

Next, we averaged decoding performance over the three classes (black trace in Figure 3, below), to assess the time-course of word class encoding, more broadly. We find that decoding performance is higher relative to word offset than word onset (mean AUC from 0-1000ms; word onset = 0.515; word offset = 0.52; t value = 2.7; p = 0.02). Furthermore, we find two reliable peaks in decoding performance. Relative to word onset, averaged over subjects, the peaks occur at around 110ms and 680ms. Relative to word offset, they occur at 390 ms and 600 ms.

**Figure 3:**
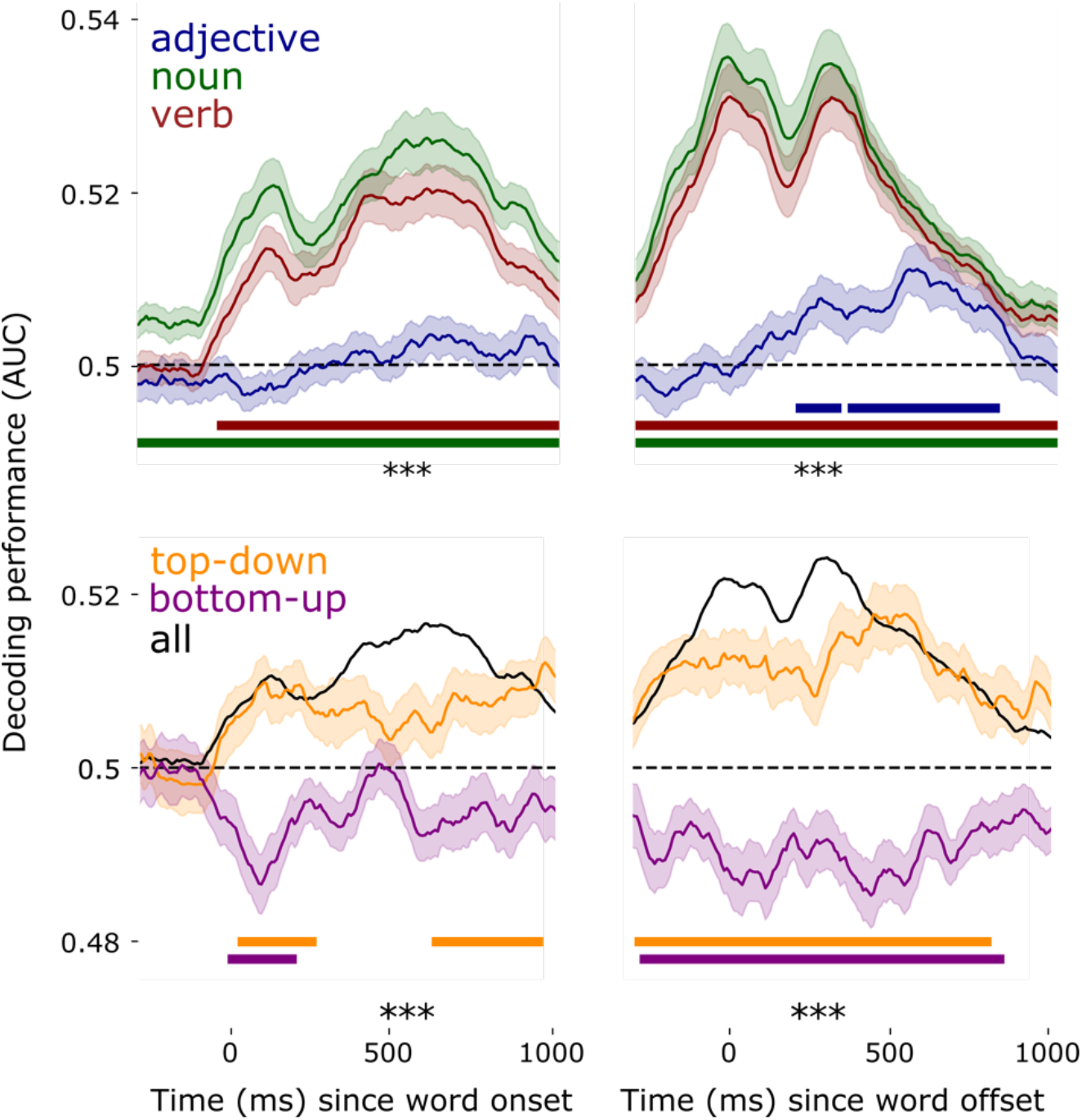
Timecourse of word class decoding. Above: Result of decoding the three word classes time-locked to word onset (left) and word offset (right). Below: Comparing bottom-up and top-down labels to the decoding model’s predictions on the mismatch trials. Horizontal lines below the timecourse represent the significant temporal clusters resulting from the permutation test. *** = p < .001. Shading represents standard error of the mean across subjects.

Overall, this first set of analyses confirms that word class is encoded in neural activity, and it is maximally decodable around 400-600 ms after word offset.

Second, and most critically for the aims of the current study, we assessed whether the neural representation of word class, in the way it is encoded during continuous speech, follows a “bottom-up → top-down” sequence of representation, or whether word class complies with top-down context directly. This is the analysis for which we plot predictions in Figure 1. We trained a logistic regression classifier on trials where the bottom-up and top-down labels of word class matched (same as above), and we examined the predictions of trials where they diverged (see Methods for details). We found significant decoding of top-down labels relative to word onset, from 30-260 ms (average t = 2.8, p = .03) and 650-1100 ms (average t = 2.7, p = .005). They were also decodable relative to word offset (−280-800 ms; average t = 3.7, p < .001). We found that decoding of bottom-up labels were significantly *worse* than chance in all cases, relative to word onset (0-190 ms; average t = -3.0, p =.02) and relative to word offset (−250-840 ms; average t = -3.1, p < .001). Our results align with Hypothesis 2 shown in Figure 1: Top-down labels are predicted above chance throughout the timecourse of reliable decoding; bottom-up labels are predicted below chance throughout. This demonstrates that the representation of word class that emerges during continuous speech processing is significantly similar to the top-down definition of word class, and significantly dissimilar to the bottom-up definition of word class.

While this is an interesting result, there is a potential confound that we needed to consider. Namely, that apparent top-down effects may actually be a confounding response to the previous word in the sequence. For example, in context, verbs are often preceded by particles like ‘to’, and nouns are often preceded by determiners like ‘the’. As such, it is possible that our classifier is learning the response to the words which systematically come before our words of interest, rather than reflecting a response to our words of interest per se. We tested this potential counter-explanation in two ways. First, we re-ran our same analysis time-locked to the onset of the preceding word. We found that neither bottom-up or top-down word class of critical w0 could not be decoded from the neural response to w-1 (all uncorrected p-values > .9, no temporal clusters found. Average effect size of top-down=0.5, average t-value=-0.14; average effect size of bottom-up=0.49, average t-value=-0.011). Second, we modified our analysis by removing all items from the train set which shared preceding words with the test set. For example, if ‘to’ preceded a verb in our test set, we removed all instances of ‘to+verb’ in our training set. This ensured that the classifier was unable to use prior word information to predict word class labels. This analysis yielded weaker but qualitatively similar results. Neither top-down or bottom-up labels were significantly decodable relative to word onset, although numerically top-down decoding was above chance (mean AUC=0.503) and bottom-up decoding was below chance (mean AUC=0.498). Time-locked to word offset, top-down labels were significantly decoded above chance from 144-672ms (average t-value=2.77; p=.005) and bottom-up labels were significantly decoded below chance from 512-880ms (average t-value=-2.5; p=0.01). In sum, both analyses provide evidence that our top-down word class decoding effect is not driven by the potential confound of sensory processing of the prior word.

Building from this, we observed that word class decoding does not decrease to chance level immediately after the word ends. Rather, we see that it remains present until around 400ms past the offset of the *next* word (Figure 4, lower right panel). This demonstrates that lexical information is long-lived, and it is maintained while subsequent words are entering into the system.

**Figure 4.**
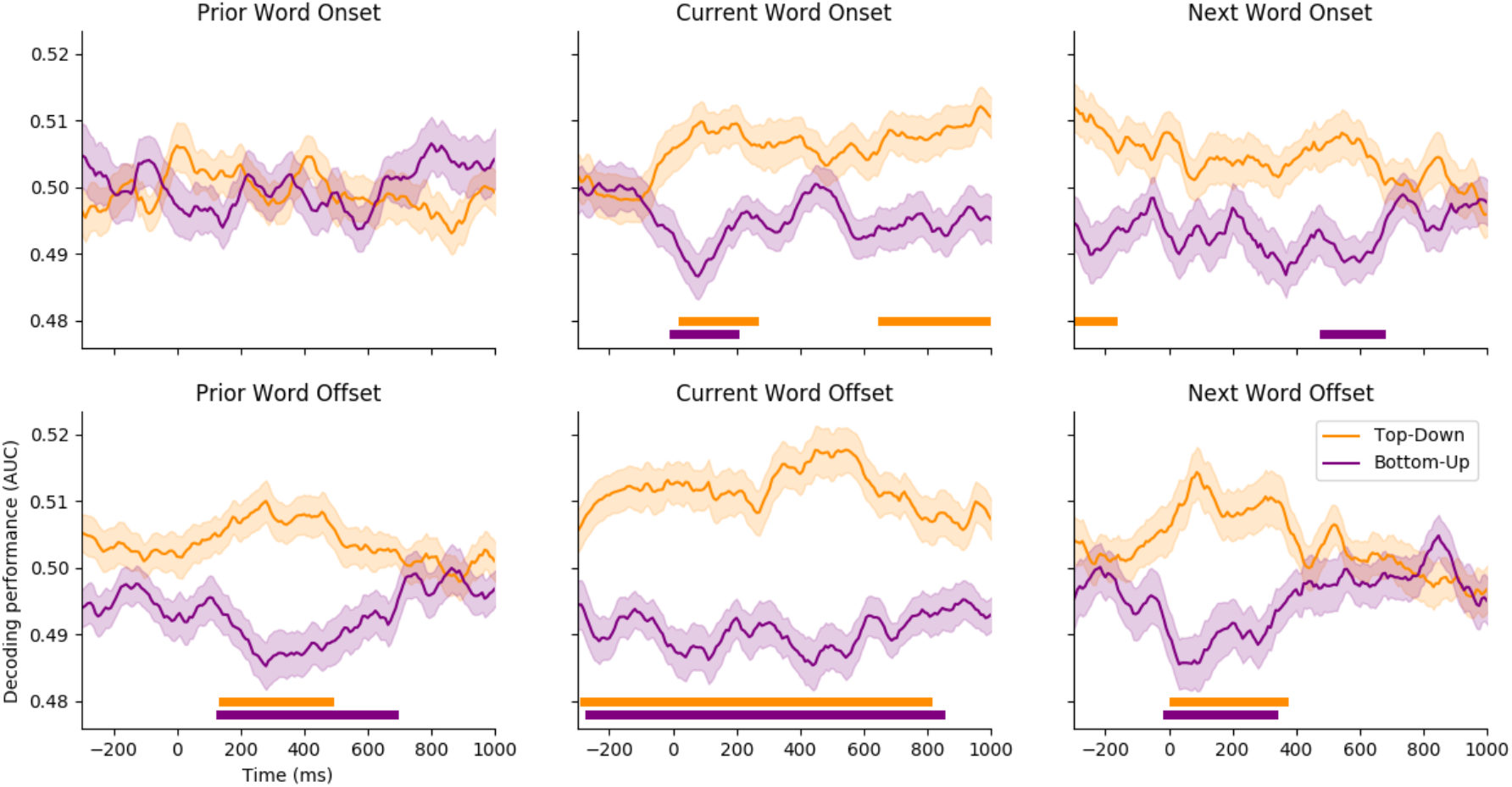
Timecourse of word class decoding at different word positions. Left: Decoding timelocked to the prior word. Middle: Decoding timelocked to the current word (recapitulation of the results in Figure 3). Right: Decoding timelocked to the subsequent word. Horizontal lines below the timecourse represent the significant temporal clusters resulting from the permutation test at p < .05. Shading represents standard error of the mean across subjects.

**Figure 5.**
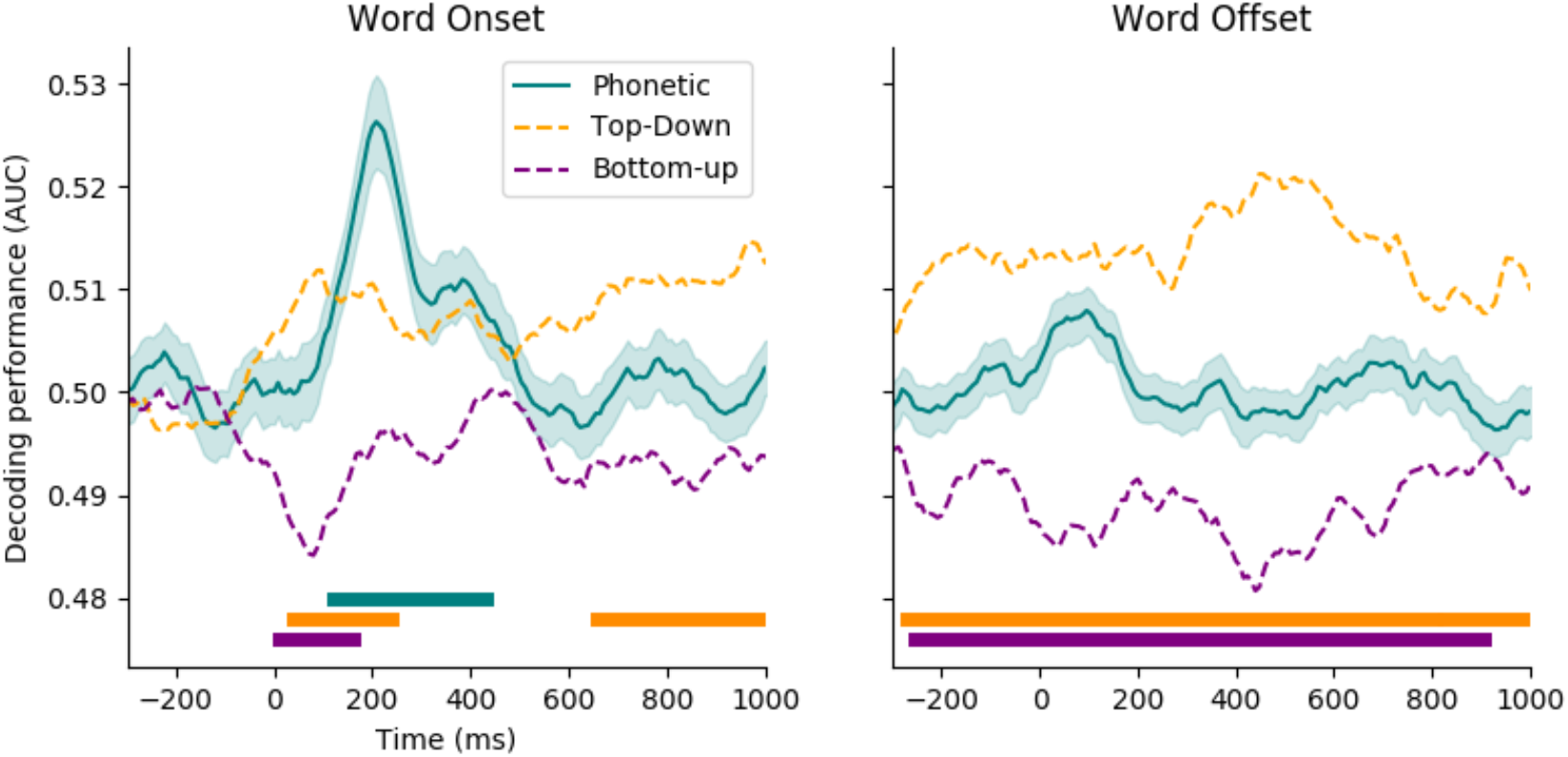
Timecourse of phonetic and word class decoding. Left: Decoding timelocked to word onset. Teal corresponds to the average decoding of voicing, fricative manner and nasal manner, of the first phoneme of the word. Orange and purple traces correspond to the part of speech decoding shown in Figure 3. Right: Same analysis time-locked to the last phoneme of the word. Teal corresponds to the average decoding of the features of the last phoneme of the word. Horizontal lines below the timecourse represent the significant temporal clusters resulting from the permutation test at p < .05. Shading represents standard error of the mean across subjects.

Given the wealth of previous work studying lexical ambiguity, which has demonstrated that bottom-up representations are accessed despite of conflicting context (Simpson, 1984; Rodd et al., 2010; Simpson and Kang, 1994), we conducted several additional analyses to test whether a bottom-up representation of word class exists in parallel to the top-down representation. One important question is whether bottom-up sensory processes are by-passed entirely, in favour of the top-down contextual representation? To evaluate this, we trained a classifier to distinguish phonetic feature contrasts of the first phoneme of our words, and the last phoneme of our words. We picked three contrasts that have been found to be strongly encoded in neural responses in previous work (Mesgarani et al., 2015; Gwilliams et al., 2022), namely: voicing, fricative manner and nasal manner. Like the analysis described above, we trained the classifier on match trials and evaluated decoding performance on the mismatch trials. This analysis yielded two findings. First, phonetic contrasts are significantly encoded in neural activity time-locked to word onset (96-488 ms; t=3.3; p=0.01). There was a numerical positive deflection of decoding the phonetic features of the final phoneme at word offset, but this did not reach significance (average t-values=0.11). In previous work we have found significant above-chance decoding for phonetic features at word offset (Gwilliams et al., 2022). We are choosing not to interpret the difference between phonetic decoding onset and offset, which could possibly be attributed to distribution of different phonetic features at different phoneme positions, or to the tendency in English to assimilate and co-articulate sounds that occur at the ends of words (Nooteboom, 1981).

The word onset result suggests that, although continuous speech processing represents word class in a top-down manner, the brain still processes bottom-up phonetic content of the word. Second, the latency of phonetic decoding proceeds word class decoding: The rise of word class decoding begins at around 0ms relative to onset, whereas it begins at around 100 ms for phonetic features, leading to a peak time of 96ms for top-down word class and 208ms for phonetic decoding. This suggests that sensory information of the word is encoded by the brain in parallel to word class information, and further confirms that bottom-up processes are not the means by which that word is initially processed, because sensory information is encoded later in time.

Building on this analysis, we sought to investigate the spatial patterns that encode word class and compare them to the spatial patterns that encode lower-level phonetic information. The subtraction between fricative and voicing evoked responses served as our phonetic contrast, and between noun and verb served as our word class contrast (Figure 4). We ran a one-sample permutation cluster test on these contrast subtractions to assess whether it significantly differed from zero (see Methods for details). In line with our decoding analysis, we found that the phonetic contrast peaked significantly later than the word class contrast. Phonetics was significant from 150-270 ms (average t-value over significant sensors = 3.3; p = .008) and the word class contrast was significant from 20-850 ms (average t = 3.1; p = .01). The responses that discriminate these word classes were largest at sensors over temporal cortex, evolving from a bilateral mid-line topography, to a more frontal topography, to a left-lateralised topography over time. The phonetic contrast was more distributed, extending from mid-line to frontal sensors bilaterally. The phonetic topography at 200 ms was significantly different from the word class topography at both 100ms (average t = 2.6; p = .014) and 200ms (average t = 2.4; p = .023. This result suggests that word class is encoded within a distinct neural population from bottom-up phonetic detail, and it is maintained for a significantly longer period of time, well into the processing of the subsequent word(s).

Another possible reason why we do not observe above-chance bottom-up decoding is that effects are masked by cases where the bottom-up and top-down labels are more equally probable. Perhaps on the trials where the bottom-up label is much more likely than the top-down label, we would be able to see evidence for bottom-up word class decoding. To test this, we extracted part of speech (POS) usage tags from four English corpora (Brown, Treebank, Conll2000 and NPSChat) and summed the number of instances of noun, verb and adjective in each. We then computed a ratio score of the top-down and bottom-up label usage based on the relative frequency of the different POS categories. We split the critical test trials into two groups, as a function of the median of this ratio metric, and decoded top-down and bottom-up labels separately in these two sets of trials.

The results of this analysis are presented in Figure 7. First, and most importantly, we find that the bottom-up label is not significantly predicted above chance in either of the two subsets of trials. This suggests that even in the context of strong mismatch, where the bottom-up word class is much more likely than the top-down word class, we still do not find evidence for a representation of the bottom-up word class label.

Second, we observed differences in the dynamics of the decoding between the weak and strong mismatch trials. When the top-down usage is rarer, the peak of average decoding occurs very late (∼900 ms), whereas when top-down usage is more common, average decoding peaks much earlier (∼100 ms). This could be readily interpreted under an evidence accumulation hypothesis, whereby a less common representation takes more time to derive than a more common one. It is interesting though, that in either case, the brain does not seem to first derive a bottom-up representation and then replace that with a top-down representation later in time: in all cases, a top-down representation is present from the beginning of processing.

Finally, we sought to understand whether a bottom-up representation of word class exists in parallel to the top-down representation, but we were unable to detect it because it occupies a distinct neural pattern. Using just the mismatch trials for training and testing, we built two classifiers: One decoding the bottom-up labels, and another decoding the top-down labels. Because of the very low number of trials included in the analysis, our estimates were too noisy to draw conclusions. Both top-down and bottom-up labels were decodable at chance level (top-down average AUC=0.501; bottom-up average AUC=0.502). Future work should seek to explore this observation further, by training a model on responses to words out of context (putatively boosting the bottom-up representation) and training a model on responses to those same words within context as we did here (putatively boosting the top-down representation). Such an approach would assess whether bottom-up representations exist in parallel, though occupying a different neural pattern, to the top-down representations.

## 4 Discussion

Deriving meaning from speech involves overcoming a wealth of sensory, (sub)lexical and structural ambiguities. In this study, we test how ambiguity of a word’s grammatical class (noun, verb, adjective) is resolved during natural story listening. We find that the neural representation of word class, as generated when processing continuous speech, more closely aligns with the contextually appropriate grammatical category of a word than the category most often realised by that word form. This suggests that context constrains how grammatical properties of lexical items are represented and interpreted.

Previous work investigating top-down processing has primarily tested the context of adverse listening conditions. For instance, to make comprehension more difficult, studies have adjusted the signal-to-noise ratio by adding noise on top of the speech signal (Zekveld et al., 2006; Davis et al., 2011) or degraded the quality of the speech signal (Hannemann et al., 2007; Sohoglu et al., 2012). Other studies have engineered language stimuli to include systematic ambiguity at the phonetic (Lee et al., 2012; Gwilliams et al., 2018) and lexical levels (Rodd et al., 2002, 2005), or provided top-down information in a different modality, such as hearing speech while reading text (Sohoglu et al., 2014). All these studies demonstrate, in different ways, that top-down information serves to resolve noise and ambiguity in the speech stimulus. Our work highlights that top-down processing is not only recruited when speech is particularly difficult to understand. But rather, even in ideal listening conditions, we observe clear evidence of top-down processing in service to grammatical processing.

The significant below-chance decoding we observe for bottom-up frequency-based labels of word class is interesting considering previous findings on lexical semantic ambiguity resolution. In that literature, it has been found that ‘bottom-up’ information, i.e., all possible meanings of that word, are activated initially, followed by the ‘top-down’ final context-congruent interpretation of the word (Simpson, 1984; Simpson and Kang, 1994; Rodd et al., 2010; Rodd, 2018). There are a few possible explanations for the difference between past semantic studies and our current results on grammatical processing.

First, it is possible that grammatical properties and semantic properties of lexical items interact with context in fundamentally different ways. The representational space of semantics is continuous and subject to coercion, and semantic interpretation can be influenced by context flexibly and creatively (Pustejovsky & Jezek, 2008; Huth et al., 2012, 2016; Gwilliams, 2020). By contrast, the grammatical properties of words are binary, and prior context serves to select grammatical interpretations deterministically (Marantz, 1997). In this way, semantic processing may operate by first activating all semantic features associated with a given word form, and then combining those features with the surrounding semantic context to yield an outcome; whereas grammatical processing may prioritize identification of the single correct interpretation, for which prior context is a much more reliable cue than word form association. This would explain why the encoding of word class reflects the final, contextually aligned, “correct” category, from the beginning of processing that lexical item. The idea that different aspects of a word (e.g., word form, semantics) are constrained by context in different ways was also put forward by Sereno et al. (2006). Here we further that proposal, by suggesting that grammatical lexical properties are another dimension of recognition that have their own way of interacting with biasing contextual information.

Second, our analysis is testing for the most dominant neural response pattern that discriminates word class in continuous speech. One possibility is that the same neural pattern encodes bottom-up labels and top-down labels, but bottom-up encoding is much weaker. In this case, our analysis may only show evidence in favour of the stronger top-down labels. In practice, however, it seems unlikely that the brain would encode two different sources of information under the same coding scheme, because this runs the risk of corrupting and mixing the two sources (Gwilliams et al., 2022). A second possibility is that top-down and bottom-up labels are encoded in two equally strong but distinct neural patterns. However, if this is the case, it does not explain why our classifier trained on match trials is unable to recover bottom-up labels when tested on mismatch trials. Given our findings, it is possible that in continuous speech, the brain selectively accesses the contextually-sensitive and lower-frequency representation of word class, and it does not represent the higher-frequency but contextually irrelevant word class label. Another possibility is that bottom-up information about word class is encoded in parallel, in a distinct and also much weaker pattern than top-down information. Our attempt to test this statistically was inconclusive due to the low number of items in our study with different top-down and bottom-up labels. Future work should explore the possibility of parallel grammatical processing by training a model on responses to words out of context (putatively boosting the bottom-up representation, because that is the only information available) and training a model on responses to those same words within context as we did here, putatively boosting the top-down representation. Such an approach would help to assess whether bottom-up and top-down grammatical representations exist in parallel, or whether top-down representations enjoy exclusive activation. In addition, it would be beneficial to conduct a replication of our study using semantic ambiguity, rather than grammatical ambiguity, to compare results directly using the same analytical protocol. We hypothesize that semantic ambiguity would yield a bottom-up-then-top-down sequence of representations, in accordance with previous work on this topic.

Interestingly, although we did not find evidence for bottom-up grammatical word class labels, we did find evidence for the processing of bottom-up acoustic phonetic input more generally. Specifically, we found that the two sources of information were decodable in parallel throughout phonetic processing, and at word onset the peak in phonetic decoding is later (∼200 ms) than the peak in grammatical class decoding (∼100 ms). This provides evidence that bottom-up sensory information is indeed available to the processing system, but it is not the means by which grammatical class labels are derived. Parallel processing of lower-level phonetic and higher-level lexical information during continuous speech processing is in line with recent MEG studies (Brodbeck et al., 2022; Heilbron et al., 2022). These studies model the variability in predictability of different sized units (e.g., phonemes and words) based on different sized context windows, and they find evidence for tracking of different sized constituents in parallel. One crucial difference between the current study and these previous studies is that here we are modelling feature representations directly, rather than the predictability of a given representational outcome. The similarity between our and these previous results, despite this crucial difference in approach, suggests that both prediction errors and representational states are processed under a hierarchy of parallel processes, therefore supporting that parallel computations are commonplace during continuous speech processing.

A major implication of our results is that, in the ecological task of story listening, the neural representation of a lexical item encompasses not just information at t0, but also information provided in previous time-steps, in the form of context. Our results aid interpretation of recent studies that use neural networks to model language processing (Qian et al., 2016; Schrimpf et al., 2021; Caucheteux and King, 2022). One key result of this previous work is that artificial models do better at predicting neural responses when they make use of longer context windows (Jain and Huth, 2018; Caucheteux and King, 2022; Goldstein et al., 2022; Caucheteux and King, 2021). Our findings anchor a concrete interpretation of this result: The boost in explanatory power is caused by incorporating contextual sentence-level information into the lexical representations that are being cross-correlated.

In a similar vein, we also found that the processing of word class of the current word is very long-lived, extending into the processing of subsequent words. We have found similar results when analysing responses at the phonetic level in previous work (Gwilliams et al., 2018; 2022). For the phonetic level of processing, we propose that prolonged information encoding serves to facilitate two operations. First, maintaining low-level phonetic detail for a long period of time provides the opportunity to re-analyse that information based on the subsequent inputs that are received. For instance, if a word initial sound is ambiguous between “b” and “p”, but then the word resolves as “parakeet”, it is advantageous to be able to integrate subsequent lexical information with the sensory information of prior speech sounds. Second, if the system has the capacity to maintain information over long time periods, this may be the means by which higher order structure is composed over time. In the case of phonetic processing, maintaining a sequence of phonemes permits generation of larger sub-lexical units such as syllables and morphemes. Here we demonstrate similar longevity in lexical processing, whereby information is maintained past the point that it has dissipated from the auditory input. We propose that this serves to permit easy repair of lexical interpretation based on subsequent input, and the online construction of constituent structures over time.

In addition, we observe that word class decoding reaches its maximum performance time-locked to word offset as compared to onset. While further work should seek to test this explicitly, one explanation is that word class is initially generated at word onset, based on prior context, and gets stronger as phonetic content of the word serves to confirm the initial context-sensitive interpretation. This is also in line with our spatial analysis (Figure 6), which shows that the topography encoding word class evolves over time. It is possible that contextual information becomes integrated with the available congruent lexical information, in order to form a grammatical representation which is consistent with both the prior and current lexical inputs.

**Figure 6:**
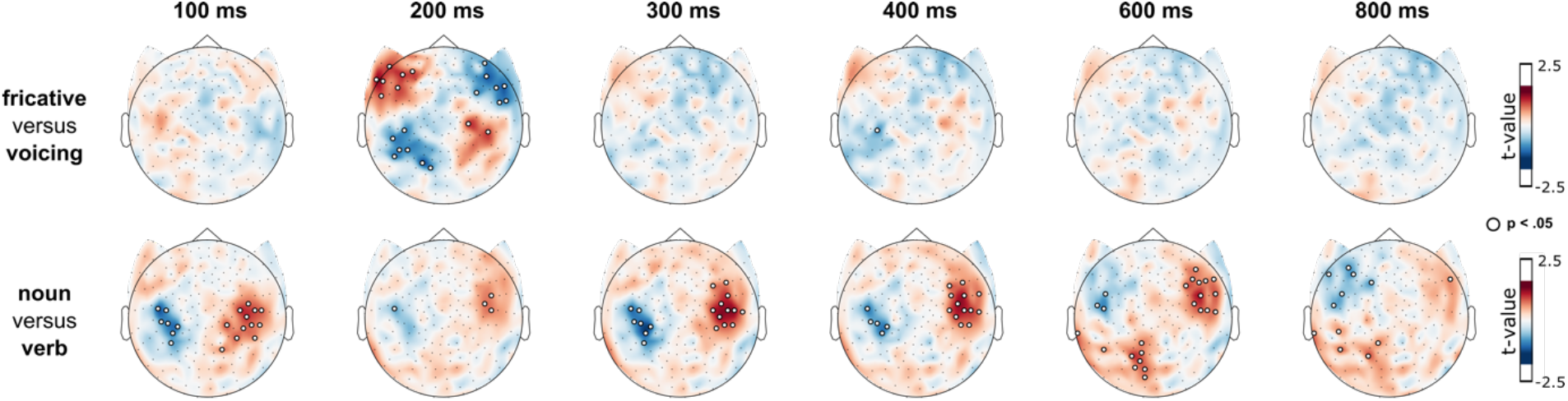
Sensor topographies of feature contrasts over time, relative to word onset. t-values over time and space for a phonetic contrast (above) and a word class contrast (below) when applying a one-sample t-test on the magnitude of the difference. Small white circles represent sensors and time-points with a significant difference between contrasts (Bonferroni corrected p < .05). Time is plotted relative to word onset.

**Figure 7.**
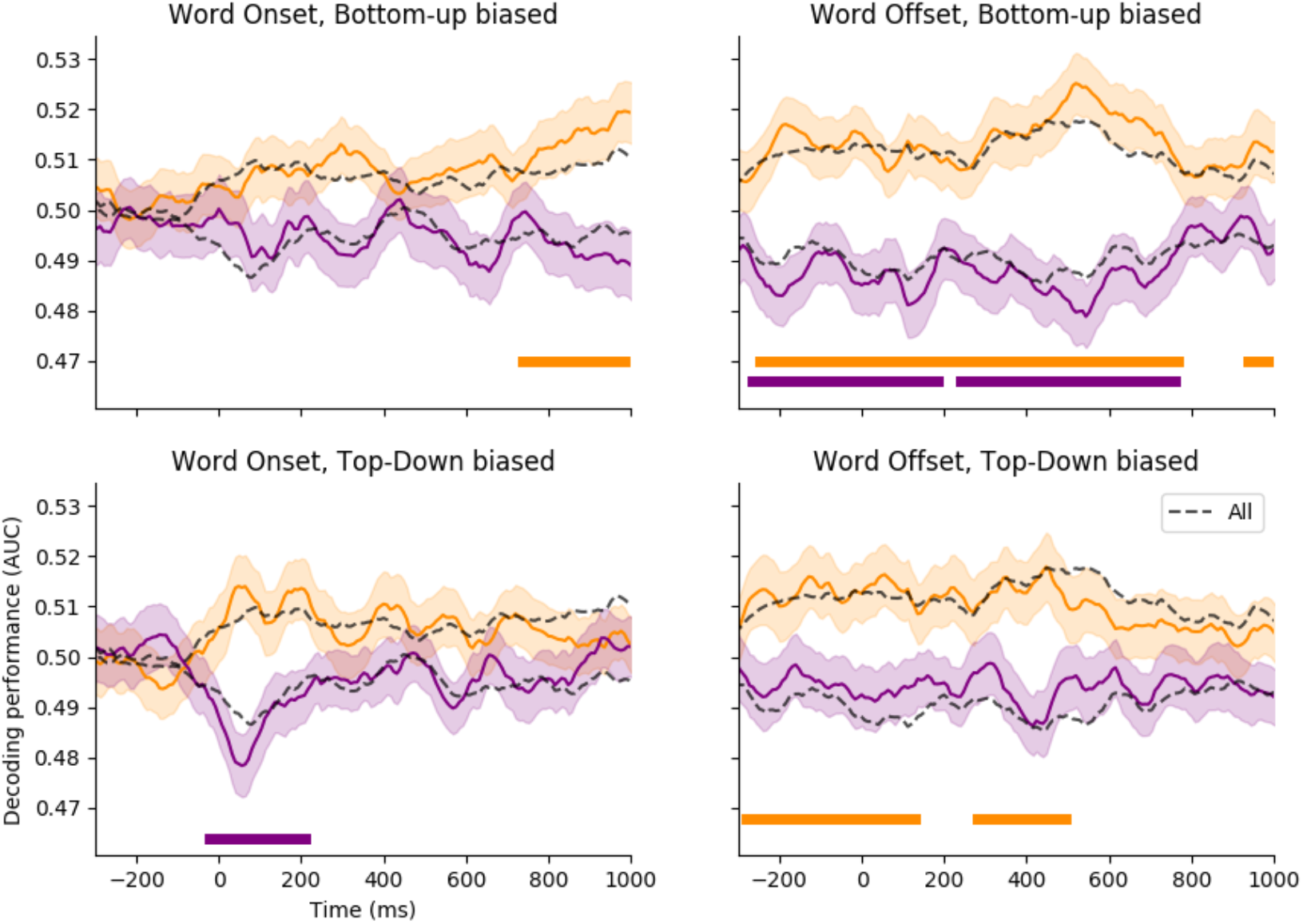
Decoding separated into trials where the bottom-up label is relatively frequent (top row) and relatively infrequent (bottom row). Orange lines correspond to top-down decoding, and purple lines correspond to bottom-up decoding. The dashed black line is the result when testing on all 202 trials, in order to allow for direct comparisons with the main result. Horizontal lines below the timecourse represent the significant temporal clusters resulting from the permutation test at p < .05. Shading represents standard error of the mean across subjects.

Taken together, our findings are inconsistent with the classic view of language processing that assumes that lower order properties of speech, closer to the sensory signal, are processed first, and serially composed into more complex abstract features over time. Here, we show that during continuous speech processing, higher-order grammatical information of speech precedes lower phonetic information, which is more akin to a reverse hierarchy (Ahissar and Hochstein, 2004; Ahissar et al., 2009). We see this explicitly in the temporal difference between word class processing (∼100 ms) and phonetic processing (∼200 ms). Concretely, this means that in the sentence “MJ looked at the stars using a…” upon hearing the word “telescope”, the brain derives that this word is a noun, before it derives that the word starts with a “t”. The reverse hierarchy theory, as put forward for visual processing, suggests that because higher order information is more robust to ambiguity in the sensory signal, it is used to guide processing and interpretation at the lower levels. We posit that a similar process is happening here for the case of speech processing: Higher order information contained in the preceding language context serves to inform interpretations at lower levels. This top-down process may aid the processing of the low-level, and generally more ambiguous, speech properties.

## Acknowledgements

This work was supported by ANR-17-EURE-0017, the Fyssen Foundation and the Bettencourt Foundation to JRK for his work at PSL and NYU Abu Dhabi Institute (G1001) awarded to AM. We thank Arianna Zuanazzi for helpful discussions on an earlier version of this manuscript.

